# Modulation of cell differentiation and growth dynamics underlie the shift from bud protection to light capture in cauline leaves

**DOI:** 10.1101/2023.12.13.571562

**Authors:** C. Le Gloanec, A. Gómez-Felipe, V. Alimchandani, E. Branchini, A-L. Routier-Kierzkowska, D. Kierzkowski

## Abstract

Plant organs have evolved into diverse shapes for specialized functions despite emerging as simple protrusions at the shoot apex. Cauline leaves serve both as photosynthetic organs and protective structures for emerging floral buds. However, the growth patterns underlying this dual function remain unknown. Here, we investigate the developmental dynamics shaping cauline leaves underlying their functional diversification from other laminar organs. We show that cauline leaves display a significant delay in overall elongation compared to rosette leaves. Using live imaging, we reveal that their functional divergence hinges on early modulation of the timing of cell differentiation and cellular growth rates. In contrast to rosette leaves and sepals, cell differentiation is delayed in cauline leaves, fostering extended proliferation, prolonged morphogenetic activity, and growth redistribution within the organ. Notably, cauline leaf growth is transiently suppressed during the early stages, keeping the leaf small and unfolded during the initiation of the first flowers. Our findings highlight the unique developmental timing of cauline leaves, underlying their shift from an early protective role to a later photosynthetic function.

**ONE SENTENCE SUMMARY:** The dual function of the cauline leaf in protection and light capture is achieved during development through a delay of cell differentiation, growth redistribution, and transient growth decrease.

## INTRODUCTION

Plant lateral organs, such as leaves and flowers, exhibit an incredible diversity of shapes that evolved to ensure their specialized functions and to adapt to their environment. For instance, leaves of different species range from simple to compound with marginal protrusions of varying sizes and shapes (Fukushima and Hasebe, 2014; Vlad et al., 2014). Leaf diversity may also be observed within individual plants as leaf morphology changes during overall plant development – a process called heteroblasty (Nikovics et al., 2006; Yu et al., 2015; Maugarny-Calès and Laufs, 2018; Tang et al., 2023). Furthermore, other flat organs, such as sepals and petals, differ greatly from leaves in their forms and sizes despite deriving from a leaf-like ancestral structure (Bowman et al., 1991; Honma and Goto, 2001; Pelaz et al., 2001).

Regardless of their diversity at maturity, all aerial organs are initiated as simple protrusions at the shoot apical meristem (Hervieux et al., 2016; Echevin et al., 2019; Silveira et al., 2022; Burian et al., 2022). All of them are also suggested to follow a common developmental program after initiation (Burko and Ori, 2013; Runions et al., 2017; Challa et al., 2021; Whitewoods et al., 2020). For instance, studies on leaf diversification revealed common growth behaviors at the global and local scales in both simple and compound leaves (Ori et al., 2007; Kierzkowski et al., 2019). Early quantitative alterations of this shared developmental mechanism are believed to account for the strong differences in final leaf shapes (Kierzkowski et al., 2019; Whitewoods et al., 2020; Wang et al., 2022). How this common program is fine-tuned to achieve specific shapes at maturity is still unclear, but involves precise molecular tuning of patterning, growth, and differentiation (Sablowski, 2015; Kierzkowski et al., 2019; Hamant and Saunders, 2020; Whitewoods et al., 2020; Tang et al., 2023).

Current evidence suggests that the balance between proliferative growth and differentiation is essential in determining the geometry of laminar organs. For instance, the rate of margin differentiation plays a significant role in shaping leaf forms, with delayed differentiation enabling extended patterning and greater leaf complexity (Donnelly et al., 1999; Shani et al., 2010; Bar and Ori, 2014; Vuolo et al., 2018; Kierzkowski et al., 2019). Local auxin maxima at the margin, facilitated by the PIN-FORMED1 (PIN1) auxin transporter, lead to the formation of protrusions such as serrations or leaflets. However, only undifferentiated cells at the leaf margin are competent to generate auxin maxima and respond to auxin by locally increasing and reorienting growth (Ori et al., 2007; Barkoulas et al., 2008; Bilsborough et al., 2011; Kasprzewska et al., 2015; Ben-Gera et al., 2016; Kierzkowski et al., 2019).

Auxin distribution at the margin has also been suggested to act globally by coordinating cellular growth orientations (i.e., growth anisotropy). In leaves, a distal auxin maximum correlates with global growth orientations converging towards the leaf tip, likely underlying organ shape tapering toward the tip (Jaeger et al., 2008; Kuchen et al., 2012). Conversely, petals exhibit a broader pattern of auxin distribution at the margin, which likely leads to the diverging growth anisotropy underlying its distally broadening shape (Green et al., 2010; Lampugnani et al., 2013; Sauret-Güeto et al., 2013). Thus, auxin plays a critical role in guiding organ formation, although the extent to which it operates locally or globally is still a matter of debate (Bilsborough et al., 2011; Kuchen et al., 2012; Kierzkowski et al., 2019; Whitewoods et al., 2020).

Understanding how plant organs acquire their shapes is essential in the context of their specific functions. For instance, rosette leaves are large organs optimized for photosynthesis, while sepals are small, protecting the internal floral organs during their development (Roeder et al., 2010; Rodriguez et al., 2014; Bielczynski et al., 2017). Cauline leaves are the last few leaves initiated during the meristem transition from the vegetative to the reproductive phase. On one hand, they are efficient photosynthetic organs, contributing to the plant’s overall energy production (Su et al., 2011). However, they also serve as protective structures for emerging floral buds during early bolting stages (Pabõn-Mora et al., 2013; Ding et al., 2023). This suggests that the development of these leaves might exhibit characteristics of both rosette leaves and laminar organs of the flower such as sepals (Hempel and Feldman, 1994; Yang and Jiao, 2016). Indeed, when floral identity genes are ectopically expressed in *Arabidopsis*, cauline leaves convert into petal-like organs, while rosette leaf development is unaffected (Krizek and Meyerowitz, 1996; Pelaz et al., 2001). Despite these intriguing features, our understanding of the developmental dynamics of cauline leaves remains unknown.

Here, we characterize the developmental mechanisms contributing to the formation of cauline leaves underlying their functional diversification from rosette leaves and sepals in the model species *Arabidopsis thaliana*. We show that cauline leaves display a strong delay in overall elongation compared to rosette leaves. Through quantitative live imaging, we demonstrate that cauline leaf functional divergence mainly relies on the early modulation of two key components: (1) the rate and distribution of cellular growth, and (2) the timing of cell differentiation. In contrast to rosette leaves and sepals, cell differentiation is strongly delayed in the cauline leaf, allowing extended cell proliferation in the leaf blade, prolonged morphogenetic activity at the margin, and growth redistribution within the developing organ. Importantly, cauline leaf growth is transiently suppressed at very early developmental stages, allowing the leaf to stay small and unfolded during the initiation of the first flowers. Overall, our results demonstrate the unique developmental trajectory of the cauline leaf that underlies the transition from its early protective role to the late photosynthetic function.\

## RESULTS

### Elongation and unfolding of the cauline leaves are delayed compared to rosette leaves

Cauline leaves exhibit both vegetative and floral features, as they are initiated from the reproductive meristem (Hempel and Feldman, 1994; Pastore et al. 2011). In contrast to rosette leaves, which have petioles for optimizing light capture, cauline leaves lack petioles but efficiently perform photosynthesis as they are located on the stem, so have direct access to light without being in the shade of other leaves (Fig. 1A-B) (Ding et al., 2023). Furthermore, in their early stages, cauline leaves serve as a protective covering for the inflorescence meristem. This role is analogous to the way sepals shield developing internal organs of flowers (Fig. 1C) (Pabõn-Mora et al., 2013; Ding et al., 2023). The presence of abaxial trichomes on cauline leaves reinforces their protective function (Fig. 1C) (Karabourniotis et al., 2020; Berhin et al., 2021).

**Figure 1.**
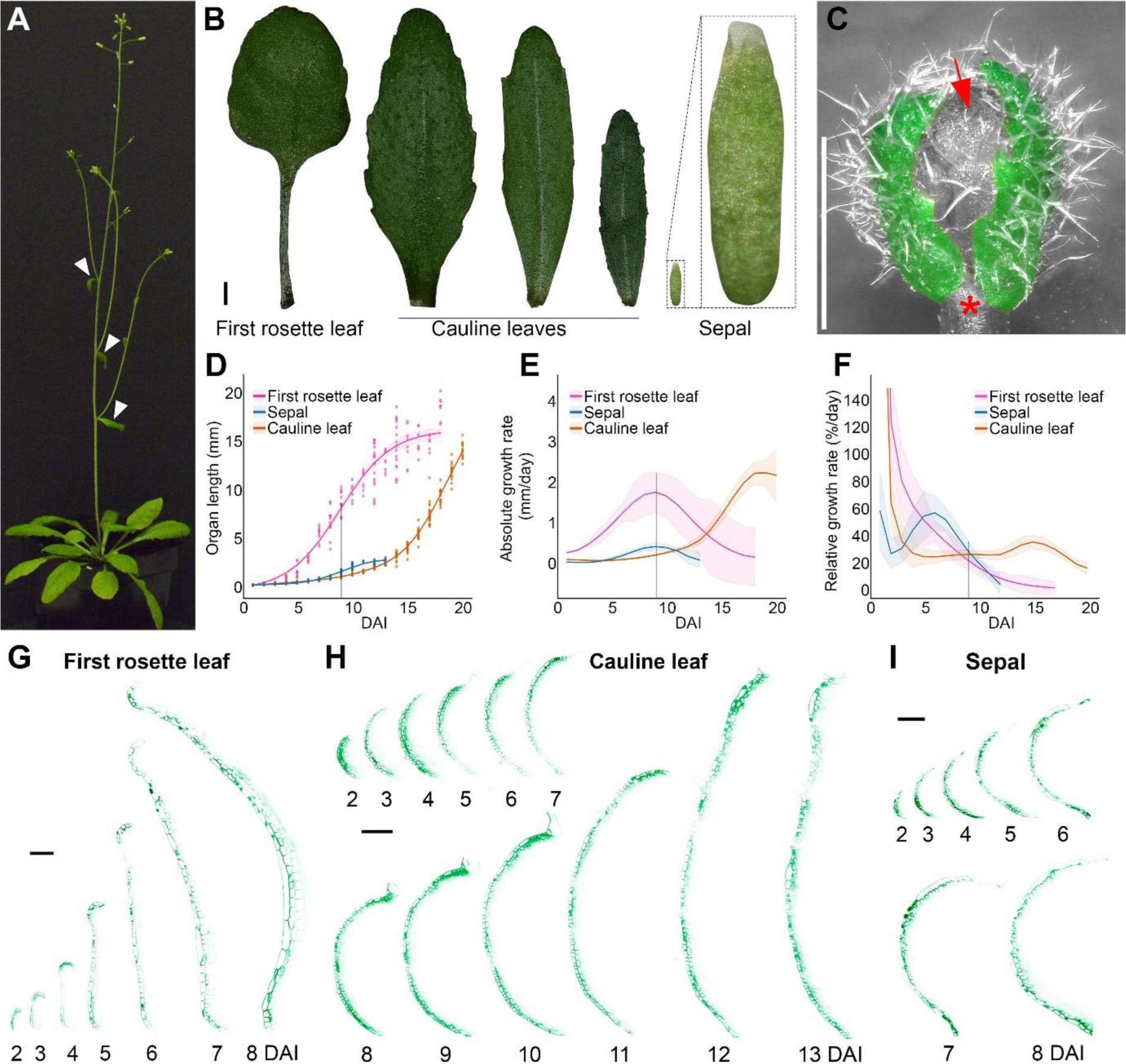
Elongation and unfolding of cauline leaves are delayed compared to rosette leaves. (A) 4-week-old *Arabidopsis thaliana* plant. Cauline leaves are indicated by white arrowheads. (B) First rosette leaf (left), cauline leaves (middle), and sepal (right) at maturity. Inset: close-up view of the sepal. (C) Cauline leaves (colored in green) cover the inflorescence meristem and developing floral buds (red arrow). Star indicates the removed cauline leaf to uncover the initiating flowers. Note the high density of trichomes on the abaxial surface. (D) Organ length plotted against time from initiation to maturity. Points represent independent samples (n=4-21 individual measurements). (E) Organ absolute elongation rate. (F) Organ relative elongation rate, showing a clear resurgence of growth in the cauline leaf at later stages. (D-F) The grey line indicates the time of peak absolute growth rate for the first rosette leaf and the sepal, at 9 DAI. (G-I) Digital, longitudinal sections located in the medial part of the developing first rosette leaf (G), cauline leaf (H), and sepal (I). DAI indicates days after primordium initiation. Scale bars: 1 mm in B-C and 100 µm in G-I. See also Fig. S1.

We supposed that the dual role of cauline leaves could be reflected in their growth dynamics. Therefore, we characterized the overall elongation of cauline leaves throughout their development and compared them with first rosette leaves and sepals. We derived two different measures of organ growth. The *absolute* elongation rate (in mm/day) simply reflects the increment of organ length over time (Fig. 1E), like earnings on interest in a saving accounts. The *relative* elongation rate (in %/day) shows how fast the organ elongates in proportion to its previous length (Fig. 1F), analog to the interest rate set by a bank. While the absolute elongation rate is more intuitive than the relative rate, the second measures growth independently from organ size and reflects better cell wall expansion.

The length of first rosette leaves evolved over time following a characteristic sigmoid shape (Fig. 1D), consistent with previous studies on leaf elongation (Cookson and Granier, 2006; Massonnet et al., 2010; Baerenfaller et al., 2015). The speed at which the first leaf elongated peaked at 9 days after initiation (Abs. rate = 1.8 mm/day ± 0.5 SE at 9 DAI) (fig 1E). This peak corresponded to the middle of the S shape, when the first leaf reached about half its final size and experienced a linear growth phase (fig 1D). Splitting our leaf measurements into petiole and blade length (Fig. S1) showed that they followed slightly different sigmoid functions, with the peak of absolute elongation occurring faster in the blade (at 8 DAI) than in the petiole (at 11 DAI), which grew slower but for a longer time (Fig. S1A). By contrast with the measurements of absolute elongation, the relative elongation rate was maximal at the very first days after leaf initiation and decreased rapidly afterwards, reaching around 20%/day at 9DAI (Fig. 1F and Fig. S1C). This fast decrease confirmed previous studies on rosette leaves (Kierzkowski et al., 2019; Le Gloanec et al., 2022; Harline and Roeder, 2023).

Growth dynamics in sepals showed similarities with the first rosette leaves, with maximal values of relative elongation rate in the days just after initiation, followed by a fast decrease (Fig. 1F). Sepals experienced their peak of absolute elongation at 9 DAI, like the first leaves, although with a much lower rate (0.42 day/mm ± 0.09 SE) (Fig. 1E), which explains the smaller length of sepals at maturity (2.6 mm ± 0.2 SE). Ultimately, the low peak of absolute elongation in sepals is due to the slower relative elongation before 9 DAI, preventing the sepal from cumulating length early on (Fig.1D).

Curiously, cauline leaves displayed a very different growth trajectory. Cauline leaves maintained a smaller size than sepals until 13 DAI (Fig.1D), due to a very low absolute elongation rate (Fig.1E). This was caused by a sharp decrease in relative elongation just after initiation and maintenance of a low relative growth rate (around 25%/day) between 4DAI and 12 DAI (Fig.1F). The first leaves and sepals reached their final size around 13 DAI, at which point they ceased growing both in absolute and relative terms. By contrast, around 13 DAI, the cauline leaf increased it growth rates, reaching a peak in relative elongation rate at 15 DAI (35 %/day ± 5 SE), followed by a peak in absolute elongation at 19 DAI. This delayed secondary growth acceleration compensated for early slow growth, allowing cauline leaves to reach the size of first rosette leaves by 21 DAI (Fig. 1D).

This transition in growth dynamics may reflect the dual function of the cauline leaf, starting with an early protective role followed by acquisition of photosynthetic competency. If this is true, the geometry of the cauline leaves at early developmental stages should resemble that observed in sepal. To test this hypothesis, we monitored the early primordia development using time-lapse imaging starting from 2 days after initiation, when all of them have comparable shape and size (refer to the ‘Materials and methods’ section for details) (Fig. 1G-I). While the sepal remained curved over the floral bud (Fig. 1I), the first rosette leaf began unfolding around 5 DAI (Fig. 1G), at the same time as its absolute elongation rate started to increase (Fig. 1E). The cauline leaf remained curled towards the floral bud until 12-13 DAI (Fig. 1H), which also coincides with the onset of its faster absolute elongation (Fig. 1E). At their early stages of development, both the first rosette leaf and the cauline leaf could protect their respective meristems. Both organs change their curvature and potentially capture light once entering their fast phase of absolute elongation. The delayed peak of absolute elongation rate for the cauline leaf could help maintain its protective role for an extended period.

### Cauline leaves display two successive waves of growth at the cellular scale

We have shown that, similarly to the sepals, in the few days after initiation cauline leaves display slower relative elongation rates than rosette leaves. Subsequently, the relative elongation of cauline leaves accelerates (Fig. 1F), allowing them to catch up in size with rosette leaves (Fig. 1D). To further understand this divergence at the cellular scale, we used our confocal time-lapse imaging pipeline to compute cell area expansion and cell elongation (in relative terms) along the proximodistal and mediolateral organ axes for each cell in the abaxial epidermis using the MorphoGraphX software (Strauss et al., 2022).

Just after initiation (2-3 DAI), relative cellular growth in all organs was fast, with the highest growth rates registered in the first rosette leaf (Fig. 2A-D; Fig. S2). In the cauline leaf and sepal, cells located at the organ tip grew the fastest, while at this stage, cellular growth rates were more homogeneous in the first rosette leaf (Fig. 2). From 3 to 7 DAI, we have observed a progressive decrease in growth rates in all organs. However, this decrease was the most dramatic in the cauline leaf (Fig. 2A- E). During this phase, a clear basipetal (from tip to base) gradient of growth was established both in the first rosette leaf and sepal (Fig. 2A-B and D-E; Fig. S2A, C, D, and F), while in the cauline leaf, cellular growth was very low and homogenous all over the organ (Fig. 2C and D-E; Fig. S2B and E). After 7 DAI, the growth rate further decreased in the first rosette leaf and sepal (Fig. 2A-B and D-E). By contrast, the cauline leaf accelerated its growth from around 7-8 DAI, and this increase continued at least until 13 DAI (Fig. 2C-D). Interestingly, cells located at the tip of this leaf were the first to increase their growth rate, especially along the longitudinal axis of the organ (Fig. 2C-E; Fig. S2B and E). A typical basipetal growth gradient started to be visible in the cauline leaf only from around 11 DAI when cells at the leaf tip ceased expanding (Fig. 2C-E; Fig. S2B and E). Overall, these results show that cellular growth rates at early developmental stages of the cauline leaf are much slower than in the rosette leaves. As a result of this early decrease in growth, the cauline leaf transiently maintains a size in the range observed for sepals. Remarkably, while rosette leaves and sepals progressively slow down growth over their development, the cauline leaf increases its growth at later stages. Thus, two successive waves of growth in the cauline leaf seems to enable its dual function.

**Figure 2.**
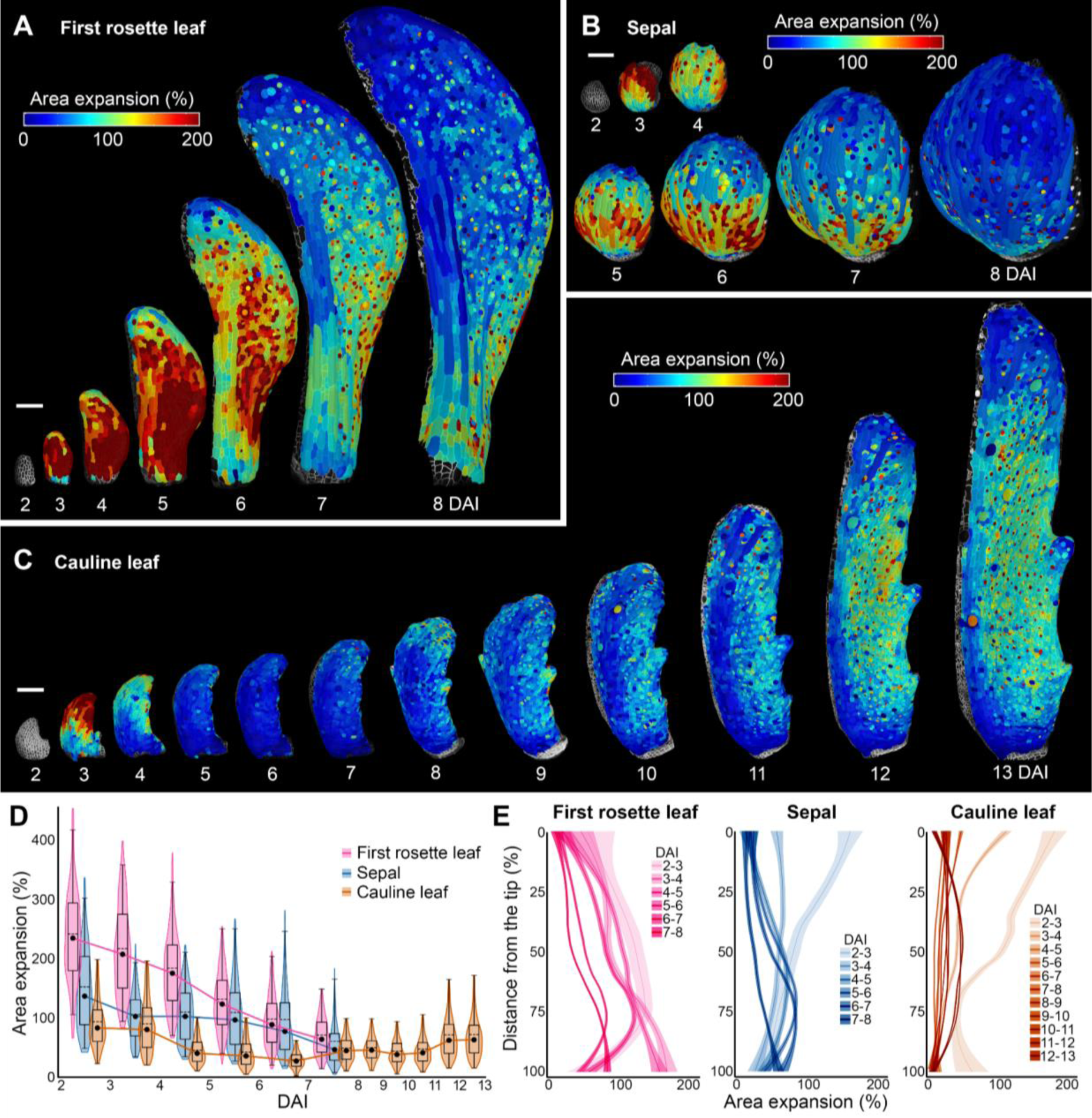
Cauline leaf displays two successive waves of growth. (A-C) Heat maps of area expansion for the *Arabidopsis thaliana* first rosette leaf (A), sepal (B), and cauline leaf (C). (D) Quantifications of area expansion for observed laminar organs. Violin plots and boxplots represent 90% of the values; mean is indicated by a dashed line, median by a black dot connected by a full line (n=810-5827 cells, three independent time-lapse series). (E) Quantifications of area expansion along the proximodistal axis of the first rosette leaf (left), sepal (middle), and cauline leaf (right). Ribbon plots display the normalized distance; the mean is represented by a line, standard deviation by the shaded area (n=58-2626 cells, based on time-lapse series shown in A to C). DAI indicates days after primordium initiation. Scale bars: 100 µm. See also Fig. S2.

### Cauline leaf maintains cell divisions during early and late growth phases

During aerial organ morphogenesis, cells located in fast-growing tissues usually divide more frequently compared to slow-growing areas (Andriankaja et al., 2012; Fox et al., 2018; Zhang et al., 2020; Le Gloanec et al., 2022; Harline and Roeder, 2023). Consistently, the basipetal growth decrease in both the first rosette leaf and sepal strongly correlated with the decrease of cell divisions from the organ tip to the base (Fig. 2A, C; Fig. 3A, C). In both organs, cell proliferation activity was first very intensive but quickly decreased at later stages (8 DAI), consistent with a progressive slow down of growth (Fig. 2A, C and D; Fig. 3A, C and D, F). By contrast, the cauline leaf retained its cell proliferative activity until at least 13 DAI, correlating with its prolonged growth phase (Fig. 2B and D; Fig. 3B and E). Surprisingly, cell divisions were also maintained in the cauline leaf during the transition through its phase of slow growth between 4 and 7 DAI (Fig. 2B and D; Fig. 3B and E). Cell divisions and differentiation are intricately linked, and a delay in differentiation often leads to prolonged proliferative activity (Vuolo et al., 2018; Werner et al., 2021; Wu et al., 2021; Gómez-Felipe et al., 2023). This trait of the maintenance of cell division may suggest that cells in the cauline leaf differentiate later than those in sepals and rosette leaves.

**Figure 3.**
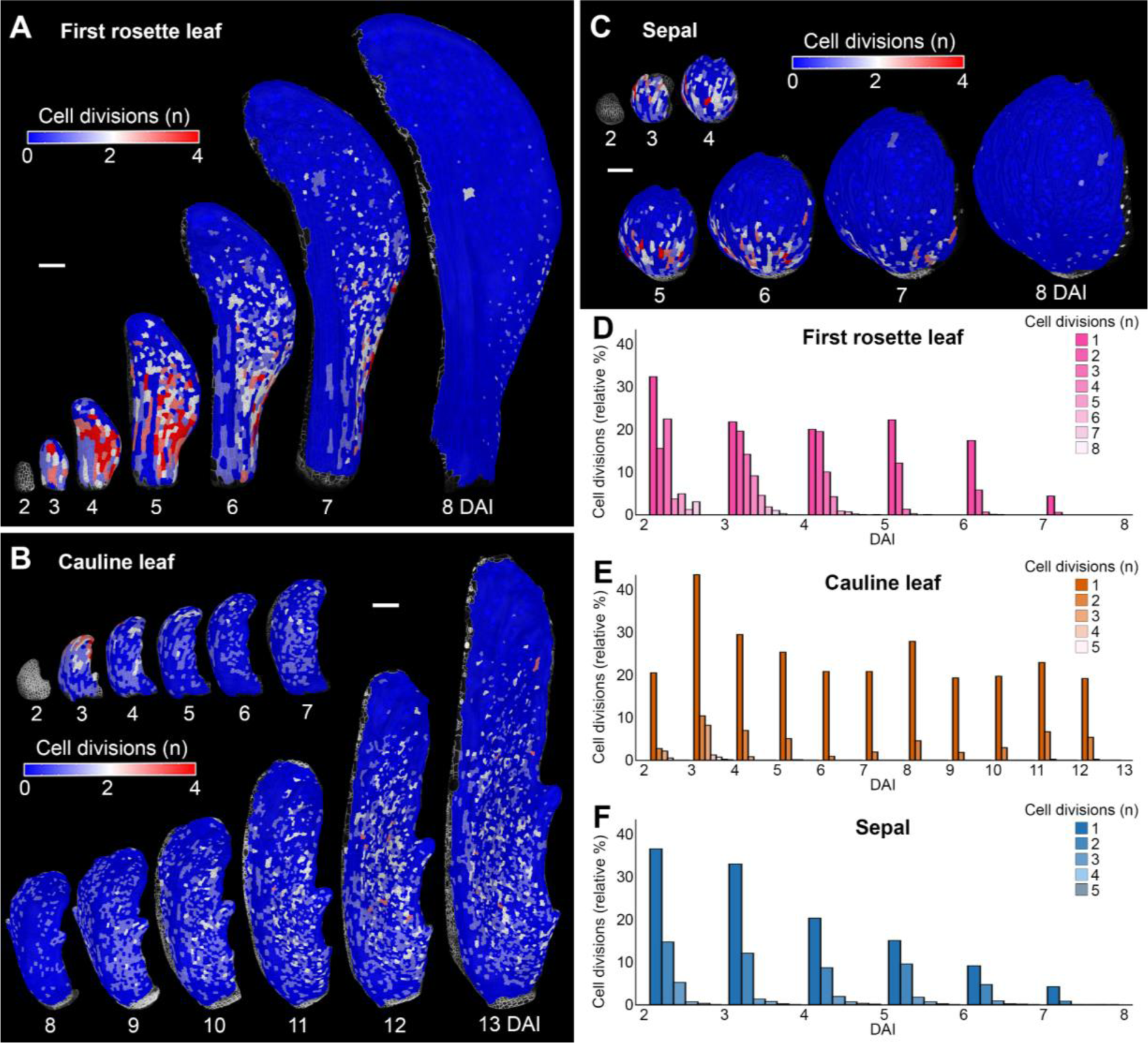
Cauline leaf maintains cell divisions during early and late growth phases. (A-C) Heat maps of cell divisions for the *Arabidopsis thaliana* first rosette leaf (A), cauline leaf (B), and sepal (C). (D-F) Quantifications of cell divisions in first rosette leaf (A), cauline leaf (B), and sepal (C). Barplots represent the relative proportion of cell divisions, normalized by the total number of cells (n=812-5828 cells, three independent time-lapse series). DAI indicates days after primordium initiation. Scale bars: 100 µm.

### Cell differentiation is delayed in the cauline leaf

To verify if cell differentiation is indeed delayed in the cauline leaf, we measured morphological markers of epidermal cell differentiation status such as cell size, stomata distribution, and pavement cell lobeyness (Andriankaja et al., 2012; Rodriguez et al., 2014; Sapala et al., 2018; Le Gloanec et al., 2022). At 8 DAI, the epidermis of first rosette leaves and sepals at 8 DAI consisted of big cells, which often developed extensive lobeyness in the leaf blade and margin or distal region of the sepal (Fig. 4A-C; Fig. S3). At this stage, stomata were already present all over the blade of the first rosette leaf and the entire abaxial epidermis of the sepal (Fig. 4D, F-G). By contrast, the cauline leaf epidermis, including the leaf margin, was mostly composed of small isodiametric cells. Only a few cells located at the tip of the first rosette leaf started to increase their size and became lobey (Fig. 4A-C; Fig. S3). Occasionally, some individual stomata could be found in the distal region (Fig. 4E and G), indicating that, at this stage, the cauline leaf is mostly undifferentiated and non-photosynthetic. Stomata spread basipetally through the epidermis only at later stages (from 10 to 13 DAI), suggesting a delay in the establishment of photosynthetic activity in this organ (Fig. 4E). Thus, organ differentiation progresses much faster in both the first rosette leaf and sepal while it is markedly delayed in the cauline leaf.

**Figure 4.**
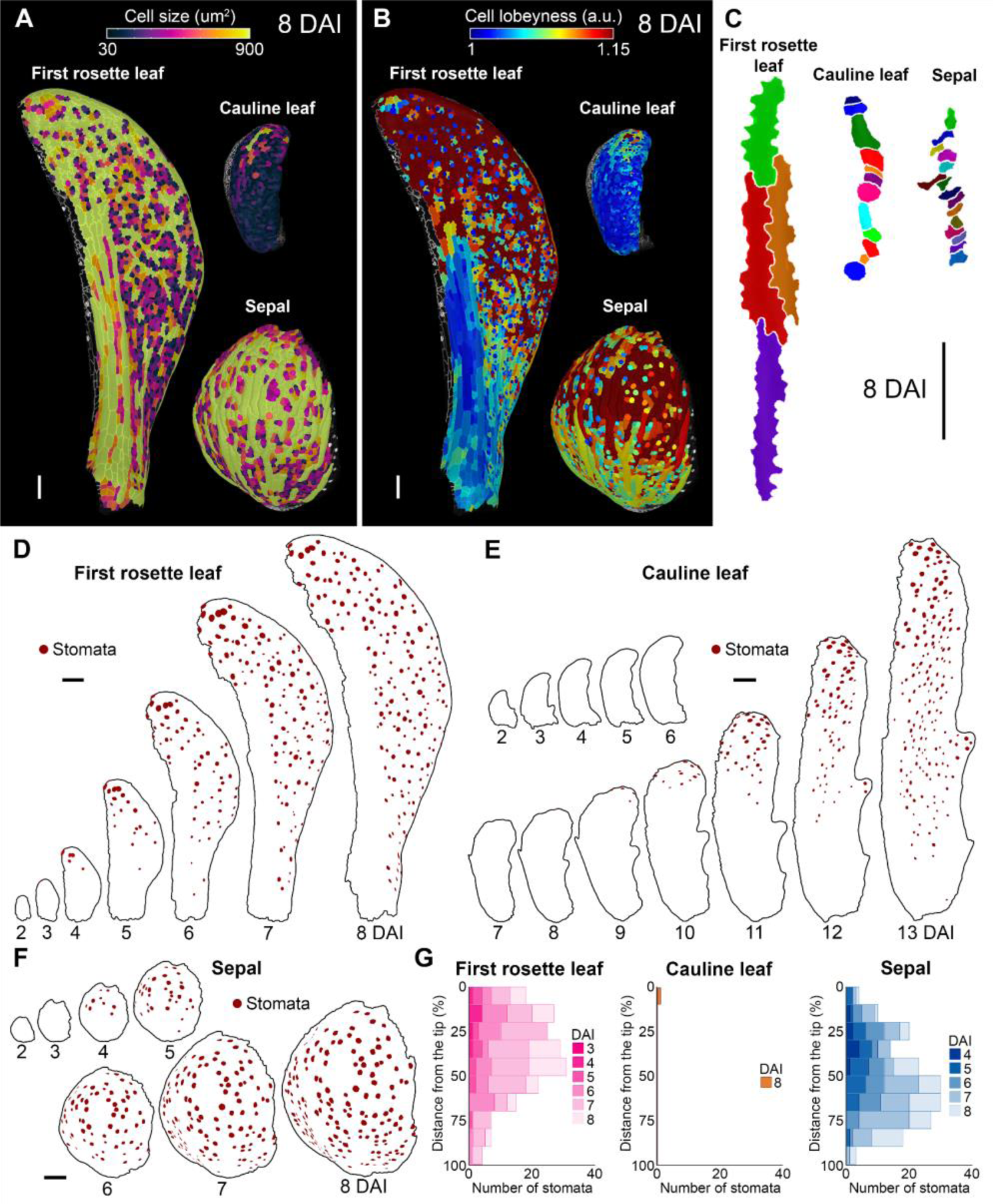
Cell differentiation is delayed in the cauline leaf. (A) Heat maps of cell size in the first rosette leaf (left), cauline leaf (top), and sepal (bottom) at 8 DAI. (B) Heat maps of cell lobeyness in the first rosette leaf (left), cauline leaf (top), and sepal (bottom) at 8 DAI. (C) Geometries of representative cells located at the distal margin of the first rosette leaf (left), cauline leaf (middle), and sepal (right) at 8 DAI. (D-F) Stomata distribution in the first rosette leaf (D), cauline leaf (E), and sepal (F). Stomata marked in brown. (G) Quantification of stomatal distribution as a function of the distance from the tip of the organ for the first rosette leaf (left), cauline leaf (middle), and sepal (right). Bar plots represent the number of stomata. (n=1-189 stomata, based on the time-lapse series shown D to F). DAI indicates days after primordium initiation. Scale bars: 100 µm. See also Fig. S3.

### Growth is redistributed in the cauline leaf

During early development, cauline leaves display a strong delay in cell differentiation compared to the sepals and rosette leaves of the same size (Fig. 4 and Fig. S3). As cell differentiation first occurs in distal portions of these organs, this may lead to an increased contribution of the cells localized at the tip of the early primordium to the final surface of the cauline leaf. Such redistribution of growth, caused by the delay of organ differentiation, has been shown to underlie the development of leaf complexity in Brassicaceae (Kierzkowski et al., 2019).

To evaluate this possibility, we computed clonal lineages that developed from cells in early primordia (2 DAI) until all organs reached comparable sizes of around 700 µm (Fig. 5A-C; Fig. S4). Already at this early developmental period, the first rosette leaf displayed smaller clones at the tip, with the marginal cells often stemming from a single cell. In contrast, the basal area grew very fast, producing elongated sectors contributing to over 35% of the organ length at the comparable size of around 700 µm (Fig. 5D, G, and J; Fig. S4B, E, and H). Conversely, cells located at the distal region of the early primordium of the cauline leaf grew the fastest and increased their size substantially compared to those at the base (Fig. 5E, H, and K; S4A and D). The contribution of the distal 20% of the primordium surface at 2 DAI increased to around 35% of the total organ surface at 10 DAI (Fig. 5E; Fig. S4G). In this respect, the sepal resembled the first rosette leaf, showcasing a greater contribution from basal cells and smaller clones at the tip (Fig. 5F, I, and L; Fig. S4C, F, and I).

**Figure 5.**
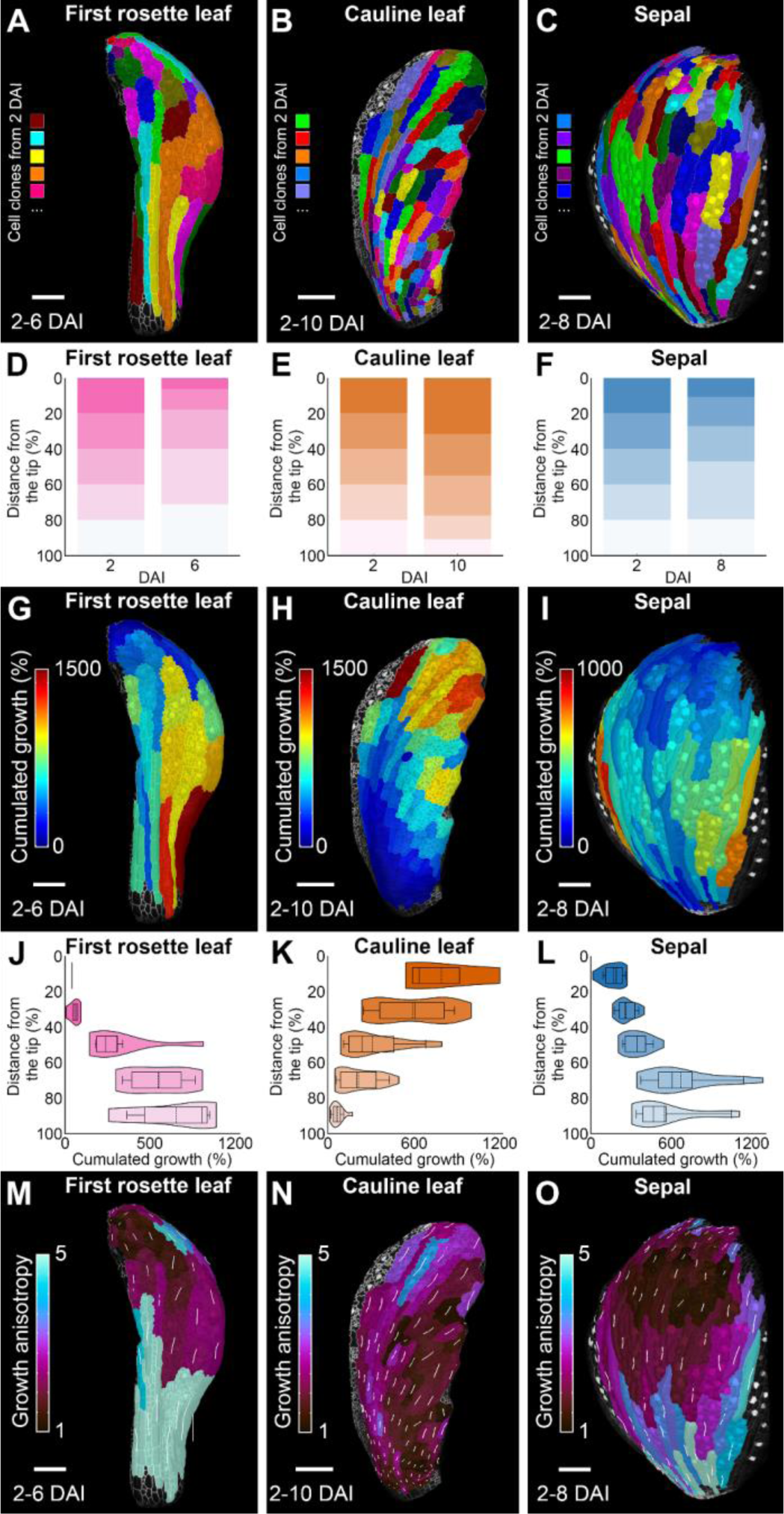
Growth is redistributed distally in the cauline leaf. (A-C) Cell lineage tracing from 2 DAI to the indicated time point (n DAI) in the first rosette leaf (A), cauline leaf (B), and sepal (C) of comparable size. (D-E) Quantification of the contribution of the clones to the length of the organ at n DAI in the first rosette leaf (D), cauline leaf (E), and sepal (F) of comparable size. (G-I) Heat maps of cumulative area expansion (from 2 to n DAI) in the first rosette leaf (G), cauline leaf (H), and sepal (I). (J-L) Quantification of the cumulative area expansion (from 2 to n DAI) in the first rosette leaf (J), cauline leaf (K), and sepal (L). (M-O) Heat maps of growth anisotropy in the first rosette leaf (M), cauline leaf (N), and sepal (O) between 2 and n DAI (comparable size). Stacked histogram represents the relative contribution of equal segments at 2 DAI to the organ length at n DAI. Violin plots and boxplots represent 90% of the values; mean is indicated by a dashed line, median by a line (n=3-7 clones, based on the sample shown in G to I). DAI indicates days after primordium initiation. Scale bars: 100 µm. See also Fig. S3.

Interestingly, the growth anisotropy of the clones in the upper half of the first rosette leaf tended to converge toward its tip (Fig. 5N). On the other hand, sectors in the cauline leaf tended to diverge from the main axis of the organ, pointing toward the distal edges of the leaf (Fig. 5N). To a lesser extent, we also observed divergent growth polarities in the sepal, but these sectors were smaller than those in the cauline leaf as they differentiated and ceased their growth early (Fig. 5M). This divergent growth anisotropy of clones in the cauline leaf may result from the maintained morphogenetic activity of its non-differentiated margin, which could act to reorient growth, as previously observed in the petal (Lampugnani et al., 2013; Sauret-Güeto et al., 2013).

### Auxin patterning mirrors the differential development of the cauline leaf

Auxin is a key regulator of cell growth and differentiation (Benková et al., 2003; Ding and Friml, 2010; Di Mambro et al., 2017; Kierzkowski et al., 2019). For instance, auxin maxima at the margin of the leaf and petal are known to locally coordinate cellular growth rates and anisotropy (Barkoulas et al., 2008; Sauret-Güeto et al., 2013; Fox et al., 2018; Kierzkowski et al., 2019; Zhang et al., 2020). To investigate if auxin patterning could underlie the differential growth observed in cauline leaves, we first monitored auxin responsiveness during organ development using the *DR5v2* reporter (Liao et al., 2015). At early stages (∼2 DAI), *DR5v2* signal was detected mainly at the tip of the first rosette leaf, and this localization was largely maintained at 4 DAI (Fig. 6A). At later stages (∼6 DAI), the *DR5v2* signal started to expand basally throughout the leaf epidermis of the first rosette leaf (Fig. 6A). Interestingly, after the initial tip localization at 2 DAI, *DR5v2* signal extended along the distal edges of the cauline leaf but did not spread throughout the leaf blade at later stages (Fig. 6B). In contrast to both types of leaves, *DR5v2* domain was broader in the distal margin of the sepal from its initiation (Fig. 6C). Like in the cauline leaf, auxin responsiveness was strong in the sepal margin but also spread through the sepal epidermis as in the first rosette leaf (Fig. 6C). A broad marginal distribution of the DR5v2 signal along the margins correlated with divergent growth anisotropy of the clones in both sepals and cauline leaves (Fig. 5). This suggests that auxin accumulation in this domain could locally reorient the growth away from the organ tip as previously observed in petals (Lampugnani et al., 2013; Sauret-Güeto et al., 2013). However, this effect was weaker in sepal as its tip quickly differentiated and stopped growing (Fig. 2B; Fig. 4A-C, and F-G).

**Figure 6.**
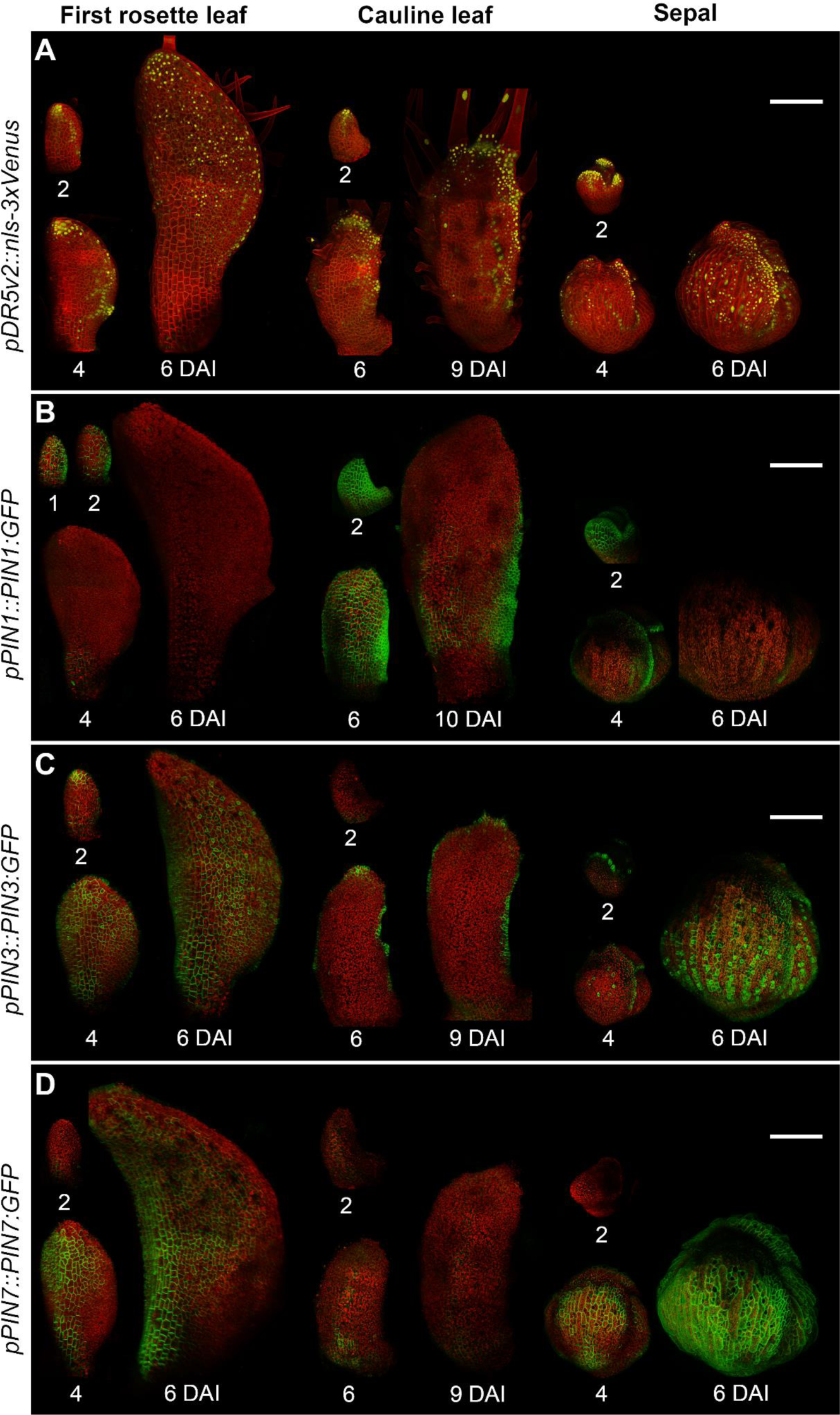
Auxin patterning mirrors the differential development of the cauline leaf. (A) Expression patterns of *pDR5v2::nls-3xVenus* in the first rosette leaf (left), cauline leaf (middle), and sepal (right). (B) Expression patterns of *pPIN1::PIN1-GFP* in the first rosette leaf (left), cauline leaf (middle), and sepal (right). (C) Expression patterns of *pPIN3::PIN3- GFP* in the first rosette leaf (left), cauline leaf (middle), and sepal (right). (D) Expression patterns of *pPIN7::PIN7-GFP* in the first rosette leaf (left), cauline leaf (middle), and sepal (right). Autofluorescence and TDT in red; Venus in yellow; GFP in green. DAI indicates days after primordium initiation. Scale bars: 100 µm.

Auxin distribution throughout the tissue is mainly regulated by auxin efflux carriers from the *PINFORMED* (*PIN*) family. Therefore, we next monitored the distribution patterns of the *GFP* fusion lines of *PIN1*, *PIN3*, and *PIN7*, which are known to be involved in leaf development (Barkoulas et al., 2008; Guenot et al., 2012; Abley et al., 2016; Mansfield et al., 2018). At early stages (1-2 DAI), *PIN1* was present throughout the epidermis of all three organs (Fig. 6B). This *PIN1* expression was quickly eliminated from the epidermis of the first rosette leaf at around 4 DAI and was restricted to the organ margin in the sepal (Fig. 6B). Interestingly, *PIN1* signal persisted throughout abaxial epidermis of the cauline leaf until at least 6 DAI, and later was still present in the proximal and lateral regions of the leaf coinciding with the initiation of the marginal serrations (Fig. 6B).

*PIN3* was expressed at the tip of the first rosette leaf and sepal from 2 DAI, while it was absent at this stage in the cauline leaf (Fig. 6C). The *PIN3* expression domain quickly (from 4 DAI) expanded basally through the epidermis of the first rosette leaf, while it was restricted to the margin of the cauline leaf (Fig. 6C). The sepal displayed an intermediate *PIN3* expression pattern: first being restricted to the organ margin at 4 DAI, then later expanding throughout the abaxial epidermis at 6 DAI (Fig. 6C). Finally, *PIN7* expression was absent at 2 DAI in all organs, and then quickly expanded through the epidermis of the first rosette leaf and sepal while it was only briefly detected in the medial and proximal regions of the cauline leaf at around 6 DAI (Fig. 6D). Altogether, we found striking differences in auxin patterning that could underlie the differential growth and differentiation patterns between these three laminar organs.

## DISCUSSION

In this study, we performed a detailed cell-level analysis of development in the cauline leaf and uncovered what sets it apart from other aerial organs, such as rosette leaves and sepals. We revealed that two main factors drive its functional distinction: (1) the timing of cell differentiation and (2) the speed and distribution of cellular growth. More specifically, cauline leaves experience a strong delay in cell maturation (Fig. 3; Fig. 7). Consequently, they display an extended period of proliferative activity and a global redistribution in growth towards the more distal parts of the organ (Fig. 2; Fig. 4; Fig. 5; Fig. 7). Remarkably, cauline leaves undergo a transient phase of slow expansion followed by a resumption of rapid growth (Fig.1; Fig. 2).

**Figure 7.**
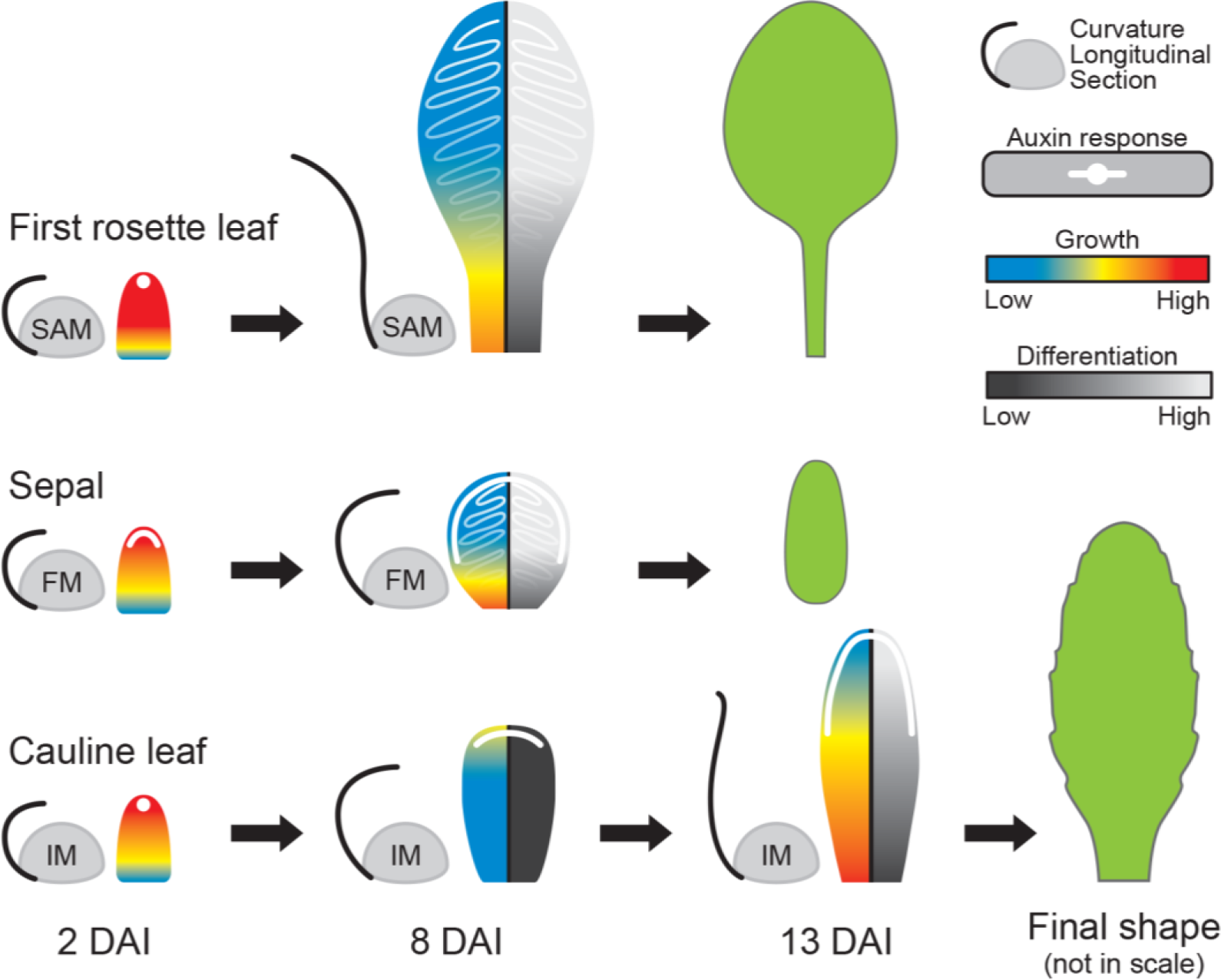
Quantitative modulation of growth and differentiation underlies cauline leaf development. Schematic representation of *A*. *thaliana* first rosette leaf, cauline leaf, and sepal shapes. Growth rates are depicted through blue-red gradients and differentiation rates by black-grey gradients, while the white color indicates the temporal distribution of auxin at specific developmental time points. All organs initiate as finger-shaped, fast-growing primordia that are curled toward meristems. In the first rosette leaf, growth and differentiation gradients are rapidly established, accompanied by auxin redistribution throughout the epidermis, and the leaf unfolds quickly. Growth and differentiation gradients in the sepal also rapidly progress; auxin is redistributed throughout the leaf margin and the epidermis, and the organ remains curled toward the flower. In contrast, the cauline leaf undergoes a transient slow-growing phase when cells remain undifferentiated, and auxin is retained at its distal margin. During this slow-growing phase, the cauline leaf stays curled toward the inflorescence. At late stages, growth rates increase again, gradients of growth and differentiation start to progress, and the leaf unfolds. DAI indicates days after primordium initiation.

Tuning cell differentiation is known to be critical for shaping plant organs (Efroni et al., 2010; Rodriguez et al., 2014; Alvarez et al., 2016). For example, delaying the transition from cell proliferation to cell maturation by class I Knotted-like Homeobox (*KNOXI*) genes has been shown to be a key component allowing the development of complex leaf shape (Hay and Tsiantis, 2006; Barth et al., 2009; Nakayama et al., 2014; Kierzkowski et al., 2019; Wang et al., 2022). Interestingly, the expression of *KNOXI* genes leads to a redistribution of growth within the organ, resulting in an increased contribution of distal organ regions to the final organ shapes (Kierzkowski et al., 2019). A similar mechanism has been observed in floral organs such as petals, where a broad distal region develops only when cell differentiation is suppressed by the action of *JAGGED* (*JAG*), which downregulates cell cycle inhibitors (Schiessl et al., 2012). Consistently, the delay of cell differentiation in cauline leaf also correlates with a greater contribution of its distal region to the organ surface (Fig. 5), suggesting that delaying the basipetal progression of cell differentiation may be a general mechanism for how plant organs change their proportions.

Interestingly, the transition from juvenile to adult phases in *Arabidopsis* leaves controlled by SQUAMOSA-promoter binding protein-like (*SPL*) genes has also been found to influence leaf morphology by delaying maturation while promoting growth and cell divisions in an age-dependent manner (Schwarz et al., 2008; Xu et al., 2016; He et al., 2018; Tang et al., 2023). *SPL10* has been shown to target genes involved in the cell cycle regulation, such as *CYCLIN D3;3* and *A2;3* (*CYCD3;3* and *CYCA2;3*) (Tang et al., 2023). As the cauline leaf marks the extremity of the heteroblastic series, the regulation of *SPL*s may underlie the strong delay in cell maturation observed in this organ. Indeed, repression of *SPL10* genes results in noticeable morphological changes in the cauline leaf, converting its shape towards the rosette leaf, including the development of a broader leaf blade (Shikata et al., 2009; Huijser and Schmid, 2011).

Auxin is known to accelerate cell differentiation and growth in *Arabidopsis* rosette leaves (Challa et al., 2019; Kierzkowski et al., 2019; Zhang et al., 2020). Here, we found that faster progression of the overall cell differentiation correlates with an early basipetal spread of auxin sensing through the abaxial epidermis of both sepal and first rosette leaf (Fig. 6). This early spread was associated with the elimination of the *PIN1* auxin efflux carrier from the abaxial epidermis that was quickly replaced by the basally progressing expression of *PIN3* and *PIN7* transporters (Fig. 6). By contrast, we observed a prolonged expression of *PIN1*, and the corresponding restriction of auxin sensing to the margin of the cauline leaf (Fig. 6). This data is consistent with recent results from the gynoecium where changes in *PIN*-mediated polar auxin transport were tightly correlated with the onset of cell differentiation in the style (Gómez-Felipe et al., 2023). Thus, modulation of active polar auxin transport may be broadly involved in controlling the basipetal progression of cell differentiation through lateral organs in plants.

Our findings indicate that marginal cells in rosette leaves undergo early differentiation, while in the cauline leaf, cells remain smaller and mainly undifferentiated (Fig. 4). *PIN3* expression expanded during the early stages of the cauline leaf development and coincided with broad redistribution of *DR5v2* signal through the distal margin (Fig. 6). Such *PIN3*-mediated auxin redistribution at the margin has been suggested to coordinate growth orientations in the petal, leading to the creation of the divergent growth polarity in its distal end (Lampugnani et al., 2013; Sauret-Güeto et al., 2013). A similar mechanism seems to operate during the early development of the cauline leaf, where growth in the distal blade is oriented toward its margin (Fig. 5). However, such reorientation of growth in the cauline leaf and petal may occur only because of their general delay in cell differentiation in the epidermis. In sepals, despite broad distribution of *PIN3*, higher accumulation of DR5v2 signal, and smaller cells at the margins, the early onset of basipetal gradient of differentiation in the epidermis quickly reduces the ability to reorient growth toward the margin.

Interestingly, both petal and cauline leaves are bent toward the inside of the flower for a prolonged period and do not quickly unfold like rosette leaves (Fig. 1) (Smyth et al., 1990; Pabõn-Mora et al., 2013). The delay of differentiation combined with the distally localized growth in the early cauline leaf may help to prevent its early unfolding. While the mechanism underlying this change in curvature remains unclear, current evidence suggests that it likely involves differential growth between both organ surfaces (Zhao et al., 2020; Jiao et al., 2021; Yadav et al., 2023). Such growth asymmetry might be controlled by the differential gene expression during the establishment of organ polarity. For instance, *ARF3* or *ETTIN* marks the abaxial side of the incipient leaf primordia and was shown to regulate the elongation of the valve tissue in the gynoecium (Andres-Robin et al., 2018; Burian et al., 2022).

In contrast to rosette leaves, which exhibit a sigmoid growth curve influenced by leaf age (Cookson and Granier, 2006; Baerenfaller et al., 2015), cauline leaves display a distinct pattern: an initial rapid decline in growth rates, followed by a surprising growth resumption (Fig. 1; Fig. 2). Although genetically induced delay of cell differentiation is known to cause decrease in growth rates (Kierzkowski et al., 2019; Wang et al., 2022), the temporary slow down in cell expansion followed by an acceleration of growth has never been documented during normal development of *Arabidopsis* leaves. However, such growth dynamics is characteristic for some floral organs, such as petals and filaments of the stamen (Smyth et al., 1990; Sauret-Güeto et al., 2013; Silveira et al., 2022). The genetic factors driving late-stage acceleration in growth remain unclear but phytohormones such as auxin, jasmonate, and gibberellins are believed to be key players, especially in the growth resumption of the stamen (Nagpal et al., 2005; Reeves et al., 2012; Cecchetti et al., 2013; Huang et al., 2020; He et al., 2023). Given the genetic proximity between cauline leaves and floral organs (Krizek and Meyerowitz, 1996; Pelaz et al., 2001), it would be interesting to investigate whether similar mechanisms govern cauline leaf development to help fulfill its dual function.

Finally, growth cessation also occurs during bud dormancy when the shoot apex inhibits axillary bud development - a phenomenon known as apical dominance (Junttila and Hänninen, 2012; Bredmose and Costes, 2017). Cauline leaves are initiated before the onset of the dormancy of the axillary buds, thus molecular signals controlling apical dominance and bud dormancy may also be involved in the transient partial inhibition of growth and subsequent growth resumption in these leaves (Rinne et al., 2011; Cooke et al., 2012; Liu et al., 2016; Salem et al., 2018; Hao et al., 2019; Martignago et al., 2020).

Our quantitative study brought new insights into the crucial role of temporal dynamics in orchestrating proper morphogenesis and functional maturation in laminar organs in plants. It provides a solid framework to investigate the molecular mechanisms underlying the development of the cauline leaf, enabling its transition from protection to photosynthesis. Such an approach could offer a deeper understanding of how these organs, originating from a common ancestor, have evolved to fulfill distinct functional roles.

## MATERIAL AND METHODS

### Plant material and growth conditions

The following *Arabidopsis thaliana* transgenic lines were used in this study: *pUBQ10::myr-YFP* (Willis et al., 2016), *pUBQ10::PM-TDT* (Melnyk et al., 2015), *pDR5v2::nls-3xVenus* (Liao et al., 2015), *pPIN1::PIN1-GFP* (Heisler et al., 2005), *pPIN3::PIN3-GFP* (Žádníkova et al., 2010), and *pPIN7::PIN7-GFP* (Belteton et al., 2018). All lines are in the Col-0 background except for the DR5 reporter line, which is in the Columbia/Utrecht background. *pDR5v2::nls-3xVenus* was crossed with *pUBQ10::PM-TDT* and observed in the 5th generation. Seeds were stratified in the dark for two days at 4°C to synchronize germination. Plants were grown on soil in a growth chamber under long-day conditions (16h/8h light/dark period, 95 µmol.m^-2^.s^-1^) at 22 ± 1°C with 60-70% relative humidity.

### Quantification and statistical analysis

Python scripts were used to generate plots in Figures 1D, 2D, F, H, 3, and 5, and Supplemental Figures 2 and 3, while R scripts were used for Figures 2E, G, I, and 4. Violin plots represented value distributions, encompassing 90% of the data. Ribbon plots indicated the mean with shaded areas representing standard deviations. Stacked histograms assessed the relative contribution of equal segments at 2 DAI during specific time points. Barplots represented the relative proportions of each value per time point using bars.

### Organ-scale time-lapse imaging and length measurements

Plants cultivated under the above-described conditions were standardized by selecting individuals of comparable sizes. Individual first rosette leaves were excised starting from 2-day-old plants every 24 h. Cauline leaves were dissected starting from 10-day-old plants every 24 h. Three successive cauline leaves from the same plant were imaged. Abaxial sepals of the first flowers were imaged every 24 h starting from 20-days-old-plant. All images were acquired using a Keyence digital microscope model VHX-970F. The lengths of all organs were measured using Fiji following the abaxial surface from the base to the tip of the organ (Rasband et al., 1997-2018).

### Analysis of organ-scale growth

Time-lapse measurements of organ length from the developing leaves, cauline leaves, and sepals were fitted to an analytical function *L*(*t*). A sigmoid function 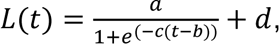, where t is the time in days, was fitted to the rosette leaf, its blade and petiole (Fig. 1D; Fig. S1) using *optimize.curve_fit* from SciPy. Splines were used to fit the length of the cauline leaf and sepal using *interpolate.BSpline* and *interpolate.splerep* with smoothing condition *s* = 0.1 from SciPy. Absolute growth rate (AGR), defined as the first derivative over time of the fitted function *L*(*t*), was calculated using the *gradient* function in NumPy (Fig. 1E). Relative growth rate (RGR) in %/day was defined as *R*(*t*) = 100 *e*^(*r(t)*−1)^, where 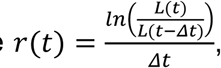 calculated from the fitted *L*(*t*) function (Fig. 1F).

### Confocal time-lapse imaging

For cauline leaf, cotyledons and older leaf primordia were removed using fine tweezers or an injection needle from two-week-old plants to expose the initiating cauline leaf primordium. The dissected plants were placed horizontally in Ø60 mm Petri dishes filled with ½ Murashige and Skoog (MS) medium, supplemented with 1.5% agar, 1% sucrose, and 0.1% Plant Protective Mixture (Plant Cell Technology). Plants were immersed in water containing 0.1% PPM for imaging. For each replicate, at least half of the abaxial leaf surface was imaged at 24 h intervals for up to 10 days. Between imaging, water was removed, and samples were cultured in vitro under standard long-day conditions in a growth chamber. The images shown in Figures 2A, 3A, and 4D and Supplemental Figures S2A, D, and S3A, D are derived from two independent overlapping series (2-7 DAI and 8-13 DAI). Time-lapse imaging of juvenile leaves and sepals was previously described (Hervieux et al., 2016; Le Gloanec et al., 2022).

All confocal imaging was performed with an LSM800 upright confocal microscope using a long-distance water-dipping objective (AP 40X/1.0; Zeiss). Excitation was performed using a diode laser with 488 nm for YFP, GFP, Venus, and 561 nm for TDT and autofluorescence. Images were collected at 500-550 nm for YFP, GFP, and Venus, 560-650 nm for TDT, and 600-700 nm for autofluorescence. Confocal stacks were acquired at 512x512 resolution and 16-bit image depth, with 0.5-2 µm distance in Z-dimension. Imaging was performed with the same confocal settings for each fluorescence marker line. For samples larger than the field of view, multiple overlapping stacks were obtained and later stitched together using MorphoGraphX (Barbier de Reuille et al., 2015; Strauss et al., 2022).

### Confocal Image analysis

Cellular growth and expression quantifications were conducted using MorphoGraphX. Stacks were processed as described previously (Le Gloanec et al., 2022). After digitally removing the trichomes when necessary, the organ surface was detected with the ‘Edge detect’ tool with a threshold of 6000-13000, followed by the ‘Edge detect angle’ (3000-6500). An initial 5 µm cube size mesh was then created and subdivided three times before projecting the membrane signal (2-5 µm).

Cell segmentation and parent attribution were performed manually. Verification of cell parenting was done using ’Check correspondence’. Lineages over multiple days were calculated as described previously and manually corrected when required (Kierzkowski et al., 2019). Stomata were identified manually based on their morphology and developmental trajectory (Le Gloanec et al., 2022).

Metrics such as area expansion, cell proliferation, growth anisotropy, lobeyness, and cell size were computed as described before (Barbier de Reuille et al., 2015; Sapala et al., 2018). Growth rates along proximodistal and mediolateral axes were determined using a custom Bezier grid, manually adjusted to follow the organ geometry at each time point closely. Proximodistal and mediolateral distances from the organ base were calculated using the ’Cell distance’ plugin, employing the ’Euclidean’ parameter.

All figures were assembled using Adobe Photoshop or Adobe Illustrator software.

## ACKNOWLEDGMENTS

We thank Binghan Wang and Jerome Burkiewicz for their assistance with data analysis and Charlotte Kirchhelle, Etienne Léveillé-Bourret, Vishwadeep Mane, Jason Reed, and Sylvia R. Silveira for suggestions and critical reading of the manuscript.

## AUTHOR CONTRIBUTIONS

C.L., A.-L.R.-K., and D.K. conceived the project and its components. C.L. and A.G.- F. designed and performed the experiments. C.L., A.G-F., and V.A. analyzed the data. C.L., E.B., and D.K. drafted the manuscript. A.G.-F., V.A., and A.-L.R.-K. revised the manuscript. A.-L.R.-K. and D.K. supervised and funded the project.

## DATA AVAILABILITY

The data used to extract growth and all scripts used to analyze data are available to download from the Open Science Framework repository (link to be provided).

## AUTHOR NOTES

The author responsible for distribution of materials integral to the findings presented in this article in accordance with the policy described in the Instructions for Authors (https://academic.oup.com/plcell/pages/General-Instructions) is Daniel Kierzkowski.

